# Overlapping enhancer activity robustly regulates *rax* expression during ocular development in zebrafish

**DOI:** 10.64898/2026.11.06.764943

**Authors:** Dwon Jordana, Nathan Clark

## Abstract

Vertebrate eye development relies on the tightly regulated yet highly dynamic spatiotemporal expression of eye field transcription factors that coordinate developmental programs required for the formation of a functional eye. The gene *rax* is among the earliest expressed eye field transcription factors and is specifically associated with ocular-fated tissue. It encodes a transcription factor that is essential for optic vesicle evagination, the precursor to the optic cup and retina. Although the developmental functions of *rax* are well characterized, the *cis*-regulatory elements that control its expression remain poorly understood.

Using zebrafish (*Danio rerio*), I identified five novel ocular enhancers and demonstrate that they drive similar spatiotemporal expression patterns as *rax* at three key stages of eye development. Furthermore, I performed an in-depth characterization of two enhancers and demonstrate that they exhibit substantial transcript-level overlap with endogenous *rax* expression during development. Additionally, through the generation of enhancer deletion lines, I further demonstrate that the gene regulatory network (GRN) controlling *rax* expression is largely robust to individual and combinatorial enhancer perturbations. However, perturbation of this regulatory network can result in *rax*-associated ocular abnormalities, including microphthalmia and anophthalmia. Together, these findings highlight the importance of studying developmental ocular enhancers to better understand the functions of noncoding genomic regions and provide insight into potential causes of unexplained congenital eye disease.

## Introduction

Ocular organogenesis requires the precise spatial and temporal expression of key genes to provide the molecular cues necessary for the morphogenetic events that underlie eye development^1^. The paired homeodomain transcription factor Rax promotes retinal identity and functions as a key component of the eye field transcription factor network^2,3^.During early development, *rax (rx3)* expression demarcates prospective ocular tissue, making it a widely used marker of eye field specification^4–6^. Furthermore, *rax* performs critical functions during optic vesicle morphogenesis and in establishing the size of ocular structures^3,7^.Additional roles for *rax* during zebrafish eye development include maintenance of the retinal progenitor cell population, establishment of the retinal pigmented epithelium, and specification of late-born ocular cell types^4,8,9^. Beyond its essential ocular functions, *rax* is also expressed in progenitor populations of the developing hypothalamus, a ventral subdivision of the diencephalon, where it is required for normal hypothalamic development and function^10,11^.

In zebrafish, *rax* transcripts are first detected in the anterior neuroectoderm at approximately 70% epiboly (8 hpf; hours post fertilization) defining the eye field^4,12^. As development progresses, *rax*-positive cells migrate laterally to form the optic vesicles^6,13^. Homozygous *rax* null mutants exhibit a recessive eyeless phenotype in which optic vesicles fail to form, resulting in anophthalmia^3^. By approximately 72 hpf, and continuing into adulthood, *rax* expression becomes restricted to the outer region of the inner nuclear layer (INL) of the retina^14,15^. Given the essential roles of *rax* at multiple stages of ocular development, mutations within this gene can lead to a variety of ocular malformations, such as anophthalmia, microphthalmia, and coloboma^16–18^.

Although mammalian *Rax* and zebrafish *rax* exhibit some differences in expression and function during the later stages of retinal development, several key features remain conserved. Both *Rax* and *rax* are among the earliest markers of the eye field and are essential for optic vesicle evagination^4,14,19,20^. Mammalian *Rax* has established roles in photoreceptor differentiation and maturation, contributes to the specification of Müller glia from late retinal progenitors, and is required in Müller glia for maintenance of retinal homeostasis in the postnatal retina^9,21,22^. *Rax* expression also coincides with the generation of late-born bipolar cells, and experimental manipulation of *Rax* expression levels alters bipolar cell abundance, supporting a role for *Rax* in bipolar cell specification^22^. Furthermore, in Xenopus, the *rax* ortholog *xrx1* is expressed within the ciliary marginal zone (CMZ), a proliferative retinal stem and progenitor cell niche that persists throughout life in fish and amphibians^23^, further demonstrating a role for *rax* in retinal progenitor populations beyond early eye development. In contrast, *rax* expression is mainly restricted to the outer edge of the inner nuclear layer, and the expression observed within photoreceptors and CMZ have been redistributed to its paralogs, *rx1* and *rx2*^14,24–26^. Expression of both orthologs, *Rax* and *rax*, has been reported within the developing INL. However, whereas *Rax* has established roles in Müller glia, *rax* expression within the INL appears to be enriched in bipolar cells, and it remains unclear whether *rax* is expressed within Müller glia^14,27^.

Though many studies have characterized the functional roles of Rax/*rax* during ocular development, comparatively little attention has been given to elucidating the cis-regulatory inputs, such as enhancers, that direct their spatiotemporal expression throughout development. Enhancers are *cis*-regulatory elements that can reside considerable distances from their target genes while directing tissue-specific patterns of gene expression^28–32^. They function as binding sites for transcription factors, including Rax, and regulate gene expression by recruiting transcription activation machinery to gene promoters in three-dimensional chromatin space^28,30,33,34^. Putative enhancers can be identified through features such as open chromatin, evolutionary conservation, and enhancer associated histone modifications, including H3K27ac^35^.

Many genes are regulated by multiple enhancers that can drive either distinct or overlapping patterns of expression^28,36^. In developmental GRNs, overlapping enhancer activities are often thought to contribute to robust transcriptional regulation by buffering against genetic and environmental perturbations^36,37^. Consequently, deletion of a single enhancer may have little or no detectable effect on gene expression or phenotype because compensatory regulatory inputs can partially maintain proper transcriptional output^36,37^. Under sensitized genetic backgrounds or following disruption of multiple enhancer inputs, however, phenotypic consequences may become more apparent^36,37^. As a result, the relationship between enhancer genotype and developmental phenotype is often non-linear, with enhancer mutations exhibiting incomplete penetrance and context-dependent phenotypic outcomes^36–39^. Nevertheless, mutations within enhancer regions can produce developmental defects that resemble those caused by mutations in the genes they regulate^36,40^. Therefore, understanding enhancer function is critical for linking genomic variation to phenotypic variation and disease^38,39^.

To date, *rax* enhancers remain poorly characterized. Only three enhancers located proximal to the *Rax/xrx1* gene body have been identified to date, largely through relatively targeted and minimalistic experimental approaches^27,41,42^. Importantly, the promoter of *xrx1* appears to be capable of recapitulating endogenous *rax/Rax* expression during eye field specification and, potentially, optic vesicle evagination. For example, transgenic studies in *Xenopus* have demonstrated that the *Rax* promoter can effectively drive expression within embryonic ocular tissues^41^. However, the complete repertoire of *rax* enhancers located more distally from the gene body remains largely uncharacterized. Furthermore, the extent to which the *rax* GRN depends on enhancer-mediated transcription to maintain robust and stable expression has not been explored. Interest in understanding *rax* enhancer function is further supported by studies implicating altered *rax* regulation in the evolution of eye loss in the cave-dwelling morph of *Astyanax mexicanus*^43–45^.

Current knowledge of enhancer-mediated *Rax/ rax* regulation does not fully address these questions. In Xenopus, the UCE3 enhancer, sometimes described as the most distal conserved element within the *Rax* promoter region, has been shown to drive expression in the embryonic eye^41^. Additionally, UCE3 represents the only reported example in which the functional contribution of *Rax/rax* cis-regulatory activity during eye development has been directly examined using optic vesicle organoids. While perturbation of this element affected *Rax* expression, its effects on optic vesicle formation were not able to be determined^46^. In contrast, UCE1 and UCE2 primarily drive expression within the ciliary marginal zone^27,42^. While these studies have provided valuable insight into *Rax* regulation, they do not define how multiple enhancer inputs cooperate to regulate *rax* expression throughout development or the extent to which individual regulatory elements are required for normal ocular development.

This study aims to identify and functionally characterize novel enhancers associated with *rax* during key stages of zebrafish eye development. In addition to defining the spatial and temporal activities of these regulatory elements, this work seeks to determine how individual enhancer perturbations influence developmental outcomes and to evaluate the potential contribution of multiple enhancer inputs to the robust regulation of *rax*-dependent eye development. The generation of stable CRISPR-Cas9 enhancer deletion alleles provides an opportunity to examine how cis-regulatory variation affects ocular phenotypes and to assess the conditions under which regulatory perturbations manifest as developmental defects. Characterization of the *rax* enhancer landscape may therefore improve our understanding of the relationship between noncoding genetic variation and phenotype, particularly in cases of incomplete penetrance where regulatory redundancy may obscure the effects of individual noncoding mutations. Furthermore, these enhancers may provide useful tools for driving cell type- and tissue-specific ocular expression in future gene therapy applications.

## Results

### Genomic Criteria Identify putative *rax* Enhancers Active During Key Stages of Ocular Development

The expression of *rax* is first detected at approximately 70% epiboly (∼8 hpf), marking the eye field^4^. However, eye field specification continues through early somatogenesis, after which the eye field bifurcates and the optic vesicles begin to evaginate^13^. To identify and characterize novel putative ocular enhancers associated with *rax* during key stages of eye development, we surveyed the *rax* locus for genomic features characteristic of enhancer activity (Fig. 1A-B)^32,47^. A previous study using *rax:GFP* reporter line demonstrated detectable expression at the 1-somite stage, consistent with our observation that the earliest enhancers activity was detected between the bud and 1-somite stages^13^. Although optic vesicle morphogenesis is occurring by 5-9 somite stage, the 7-somite stage was selected for stage representation because enhancer activity was observed across most candidate enhancers at this time point^48^. In addition, 72 hpf was selected as a later developmental stage because endogenous *rax* expression is known to become restricted to the INL of the retina^14^.

**Figure 1.**
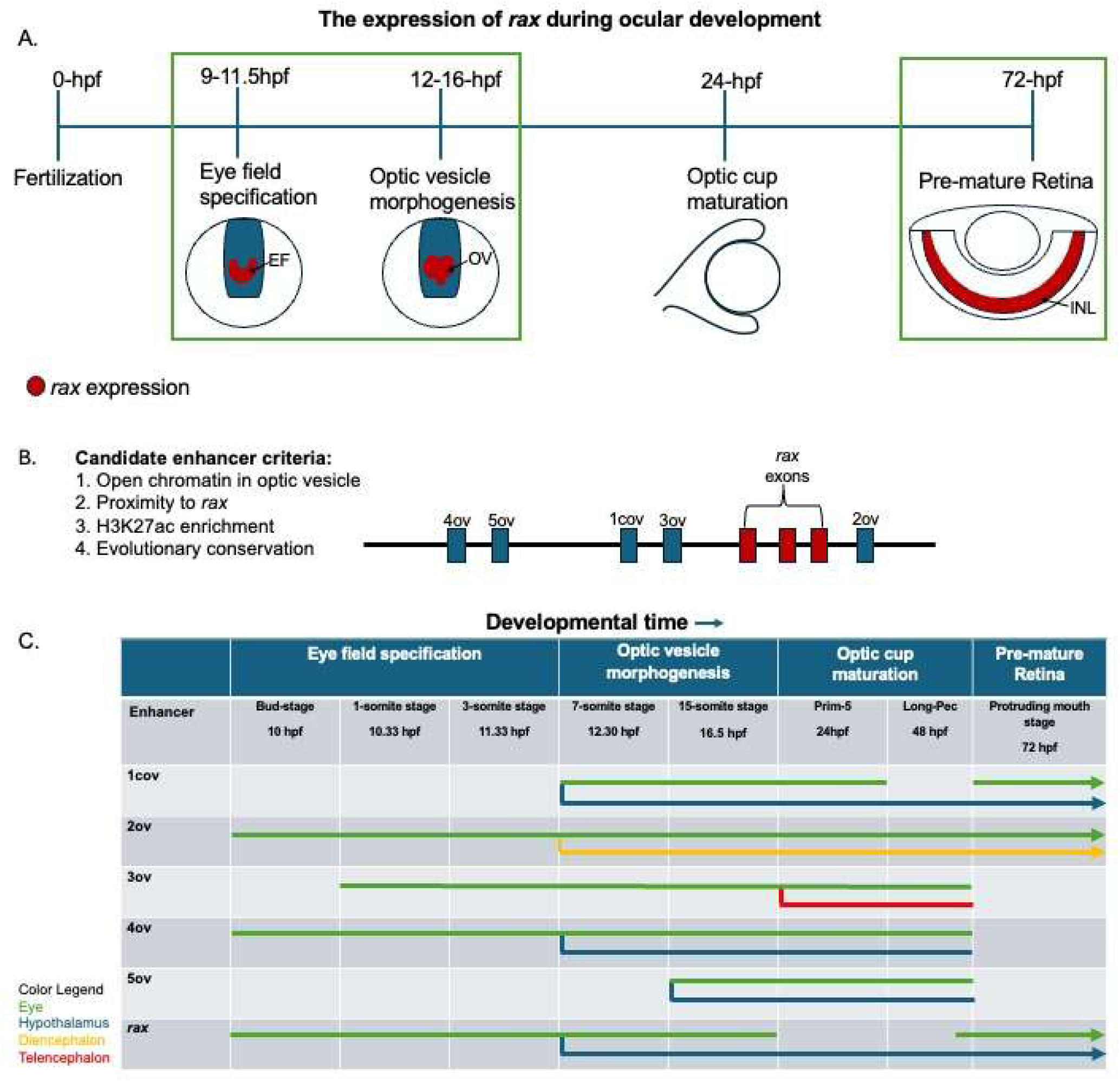
Candidate selection pipeline for putative *rax* enhancers, and overview of enhancer-driven expression patterns. (A) Zebrafish ocular development and key developmental stages relevant to *rax* expression. (B) Enhancer selection criteria, with the five highest-priority candidate enhancers shown in blue and *rax* exons shown in red. Candidate enhancer names are indicated above each region. (C) Summary of enhancer-driven transgene expression patterns during development. Green, eye; blue, hypothalamus; yellow, dorsal diencephalon; red, telencephalon. hpf: hours post fertilization, EF: eye field, OV: optic vesicle, INL: inner nuclear layer

To identify novel ocular enhancers, four genomic criteria were applied: (1) chromatin accessibility within the optic vesicle, (2) proximity within a two-megabase window surrounding *rax*, (3) enrichment for the active enhancer mark H3K27ac, and (4) evolutionary conservation (PhastCons). Application of these criteria identified several candidate regulatory regions. Five regions were selected for further analysis based on the presence of accessible chromatin within the optic vesicle. The chromosomal positions of these candidate regions relative to the *rax* gene body are depicted in Fig. 1B, and their coordinates can be found in Supplementary Fig. 1G

The spatiotemporal expression profiles of the five putative enhancers were evaluated by using a transgenic reporter assay, with each candidate region cloned upstream of a minimal promoter driving reporter expression. Putative enhancers were numbered sequentially in the order in which they were identified and designated cov (conserved optic vesicle) when sequence conservation extended to mammals or ov (optic vesicle) when identified based on open chromatin state in ocular tissues.

Reporter activity revealed distinct patterns among candidate enhancers, with the greatest variation observed in the timing of activation, whereas the tissues exhibiting enhancer-driven expression were largely conserved (Fig. 1C). The spatio-temporal expression profiles observed were as follows:

- ***1cov***-driven GFP expression was first detected at the 7-somite stage within the optic vesicle and hypothalamus (Fig. 2I). Ocular expression persisted through 24 hpf and was again detectable at 72 hpf, whereas hypothalamic expression was maintained continuously from activation through 72 hpf Fig. 3A).
- ***2ov***-driven GFP expression was first detected at approximately 10 hpf within the eye field. Following separation of the optic vesicles, expression in the dorsal diencephalon becomes distinguishable at the 7-somite stage. Ocular expression persisted from 10 hpf through 72 hpf and became progressively restricted to the anterior portion of the retina. Expression within the dorsal diencephalon also persisted throughout this period (Supplementary Fig. 1A-C).
- ***3ov***-driven expression was first observed at the 1-somite stage and through the 7-somite stage (Fig. 2A,E). Reporter activity persisted until 48hpf. In addition to ocular expression, forebrain expression was observed in the telencephalon beginning at 24 hpf.
- ***4ov***-driven GFP expression was first detected at approximately 10 hpf within the eye field. Hypothalamic expression became apparent by 24 hpf and persisted until 48 hpf (Supplementary Fig. 1D-E).
- ***5ov***-driven GFP expression was first detected at the 15-somite stage in both the developing eye and hypothalamus, with expression persisting through 48 hpf (Supplementary Fig. 1F).

**Figure 2.**
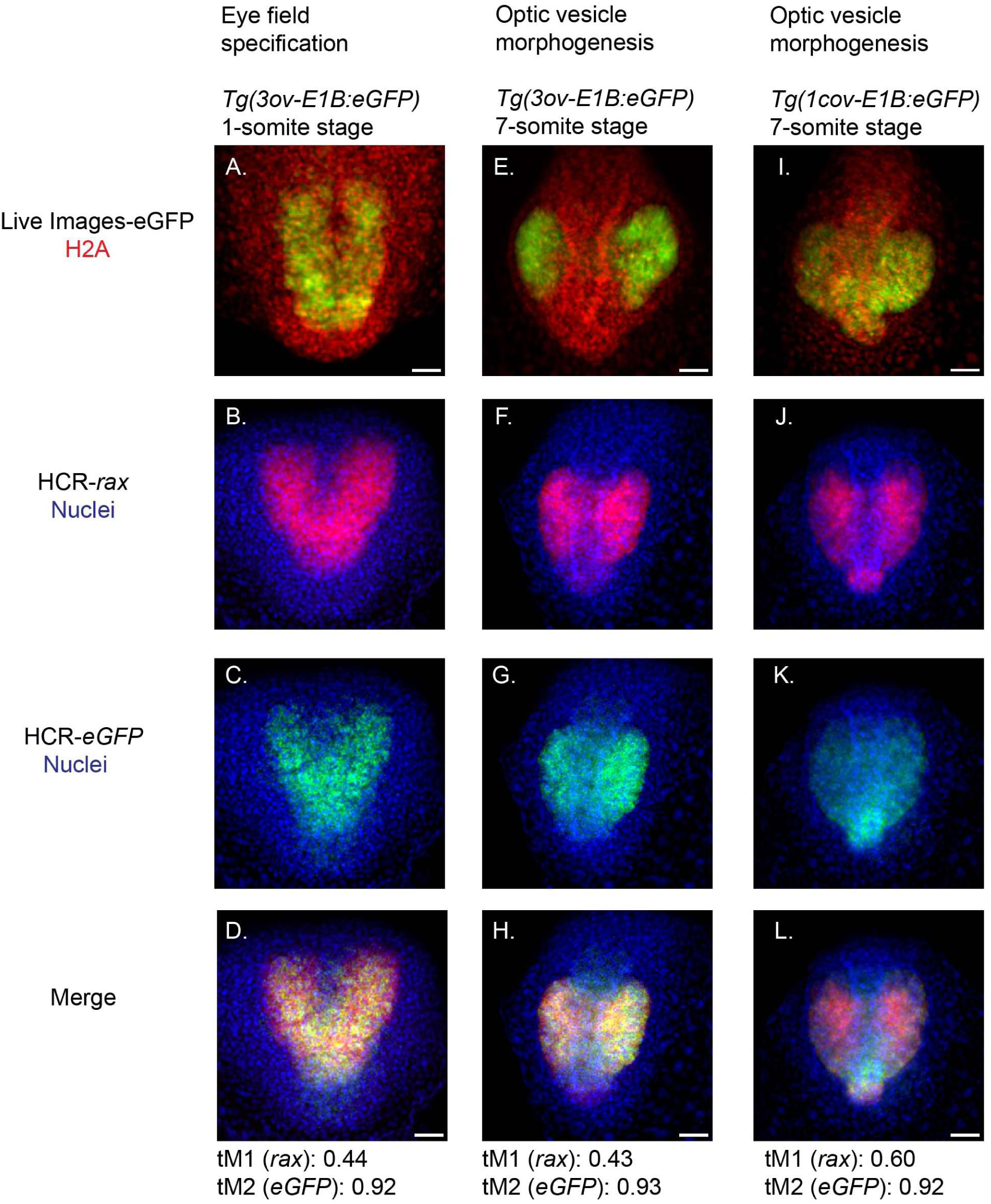
Expression of two representative *rax* enhancers during early eye development. (A) eGFP reporter driven by the *3ov* enhancer at the 1-somite stage. (B-D) HCR detection of *rax* (B) and *eGFP* (C) transcripts and corresponding merged image (D). (E) eGFP reporter driven by the *3ov* enhancer at the 7-somite stage. (F-H) HCR detection of *rax* (F) and *eGFP* (G) transcripts and corresponding merged image (H). (I) eGFP reporter driven by the 1cov enhancer at the 7-somite stage. (J-L) HCR detection of rax (J) and *eGFP* (K) transcripts and corresponding merged image (L). Thresholded Manders co-occurrence values are displayed below each merged image. Scale bars indicate 50 µm.

**Figure 3.**
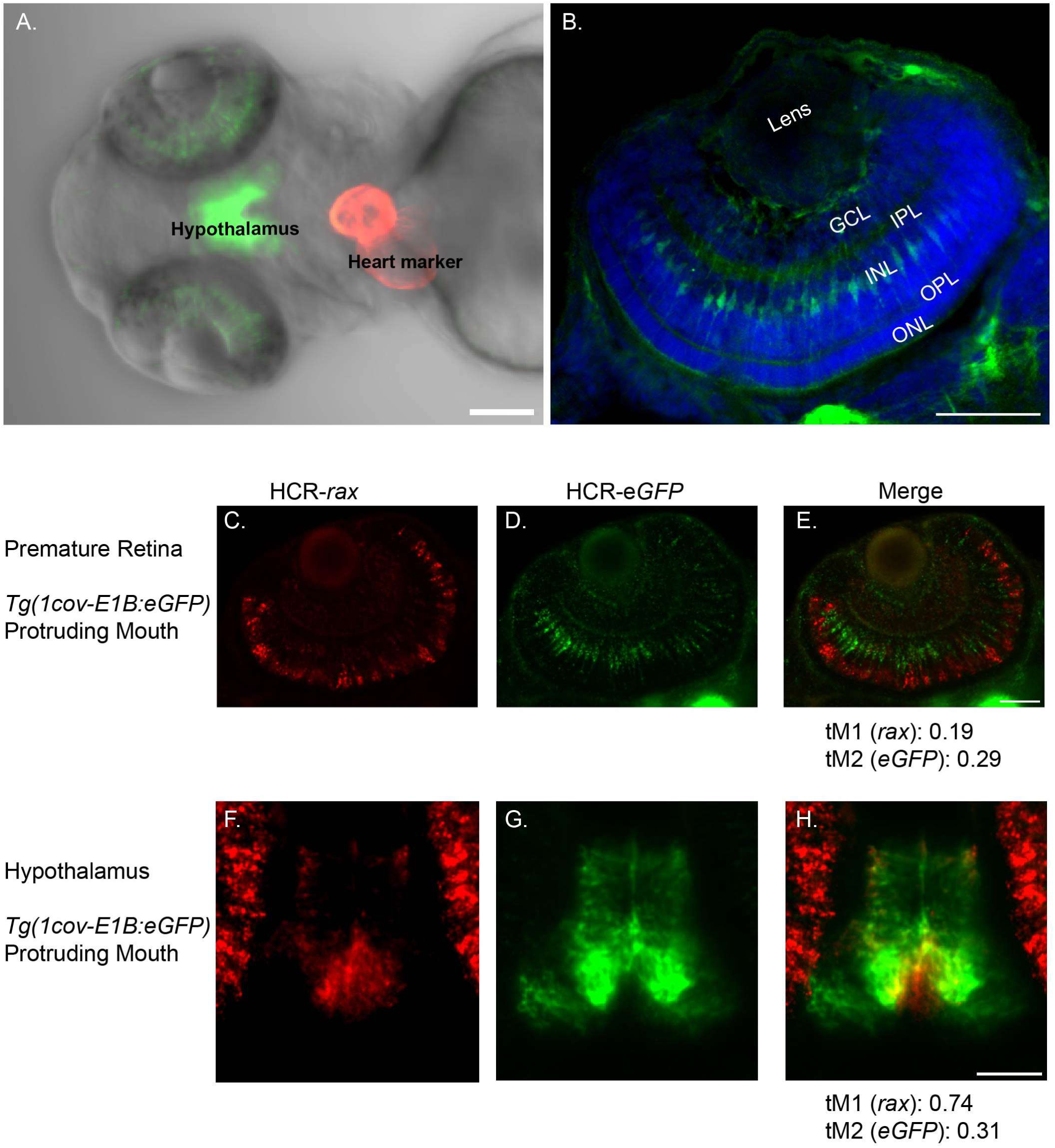
1cov drives expression that poorly recapitulates endogenous *rax* expression in the retina but not the hypothalamus. (A) Live confocal image of the *Tg(1cov-E1B: eGFP)* transgenic line at 72 hpf, showing expression within the retina and hypothalamus. (B) Retinal cryosection from *Tg(1cov-E1B: eGFP)* larvae at 72 hpf depicting radially elongated cells spanning the full thickness of the retina, with cell bodies located within the inner nuclear layer (INL). (C-E) HCR analysis of retinal tissue from *Tg(1cov-E1B: eGFP)* larvae at 72 hpf showing *rax* transcripts (C), eGFP transcripts (D), and the merged image(E). Thresholded Manders co-occurrence values are displayed below the merged image. (F-H) HCR analysis of hypothalamic tissue from *Tg(1cov-E1B: eGFP)* larvae at 72 hpf showing *rax* transcripts (F), eGFP transcripts (G), and the merged image (H). Thresholded Manders co-occurrence values are displayed below the merged image INL: Inner nuclear Layer, ONL: Outer nuclear layer, IPL, Inner plexiform layer, Outer Plexiform layer, GCL: Ganglion cell layer Scale bar in (A) 100 µm; scale bar in (B-H),50 µm.

Together, these findings identify five novel putative ocular enhancers within the *rax* locus that are active during distinct stages of ocular development. Although each enhancer displays a unique temporal and spatial expression profile, all drive expression within tissues known to express *rax*, supporting their potential roles as components of the *rax* regulatory network^14^. These expression profiles were confirmed by screening a minimum of three independent insertion events from three separate founders.

### Eye-specific enhancers drive expression within the *rax* expression domain during early development

To further investigate whether these novel ocular enhancers regulate *rax* expression, hybridization chain reaction (HCR) to the reporter gene transcript (*eGFP*) was performed at timepoints point of interest for two representative enhancers. Use of HCR on the reporter transcript provides higher temporal resolution that the eGFP protein itself, which has a long half-life. The enhancers,*1cov* and *3ov*, were selected due to the exceptional conservation of *1cov* and the close proximity of *3ov* to 1cov, as well as their ability to drive expression that best accounts for the totality of *rax* expression across the timepoints assessed in this study.

The spatiotemporal expression of the two transgenic lines was assessed by examining co-occurrence of endogenous *rax* expression with enhancer-driven reporter expression (*eGFP* transcripts) across pertinent developmental stages.

### Eye field Specification

Expression driven by the *3ov* enhancer was first detected at the 1-somite stage within the anterior neuroectoderm, corresponding to the location of the presumptive eye field at this developmental stage (Fig. 2A). Thresholded Manders overlap coefficients were used to quantify co-occurrence between *3ov*-driven eGFP and *rax* transcripts (Fig. 2B-D). The *3ov* expression domain overlapped with 44% of the endogenous *rax* expression domain (tM1 = 0.44 ± 0.05), whereas 92% of the *3ov* expression domain overlapped with endogenous *rax* transcripts (tM2 = 0.92 ± 0.02) (Fig. 2D). To assess the specificity of this interaction, a negative control was established by rotating the eGFP channel 90° clockwise. Under these conditions, the 3ov transcript overlapped with 20% of endogenous *rax* expression (tM1 = 0.20 ± 0.13), while *rax* expression overlapped with 48% of 3ov-driven transcripts (tM2 = 0.48 ± 0.24). As a positive control, *six3b*, an eye field transcription factor with overlapping expression in the early eye field, exhibited strong overlap with *rax*, although the degree of co-occurrence was lower than that observed for eGFP (Fig. 2D; Supplementary Fig. 2A-C).

### Optic vesicle morphogenesis

During optic vesicle morphogenesis, both enhancers examined in detail drove expression within known *rax* expression domains, including the optic vesicles and hypothalamus. At this stage, *3ov* drove expression exclusively within the optic vesicles (Fig. 2E). Transcript-level analysis demonstrated substantial overlap between *3ov*-driven *eGFP* and endogenous *rax* expression (Fig. 2F-H). The transgene expression domain accounted for 43% of the endogenous *rax* domain (tM1 = 0.43 ± 0.05), whereas *rax* expression overlapped with 93% of the *3ov*-driven eGFP domain (tM2 = 0.93 ± 0.09) (Fig. 2H). In contrast, negative control analysis demonstrated that 25% of endogenous rax expression overlapped with the 3ov transcript (tM1 = 0.25 ± 0.06), while 66% of the 3ov transcript overlapped with endogenous rax expression (tM2 = 0.66 ± 0.10). The positive control, *six3b*, exhibited substantial co-occurrence with *rax*, supporting the specificity of the observed enhancer activity, which more closely matched the *rax* expression domain than *six3b* (Fig. 2H; Supplementary Fig. 2D-F).

For 1cov, detectable eGFP protein expression was observed within both the optic vesicles and hypothalamus (Fig. 2I). Transcript co-occurrence analysis similarly revealed substantial overlap between enhancer-driven expression and endogenous *rax* transcripts, with the 1cov transgene overlapping 60% of the *rax* expression domain (tM1 = 0.60 ± 0.10) and *rax* expression overlapping 92% of the eGFP-positive domain (tM2 = 0.92 ± 0.03) (Fig. 2J-L). By comparison, the negative control exhibited weaker overlap between the 1cov transcript and endogenous *rax* expression, with 28% of the *rax* signal and 40% of the 1cov signal residing within the shared domain (tM1 = 0.28 ± 0.14; tM2 = 0.40 ± 0.18). While expression of six3b demonstrated strong overlap with *rax*, however not as strong as *eGFP* (Fig. 2L; Supplementary Fig. 2G-I).

To provide biological context for the observed co-occurrence between enhancer-driven eGFP and endogenous *rax* transcripts, positive (*six3b*) and negative (eGFP channel rotated 90° clockwise) control analyses were performed. Co-occurrence between *rax* and the eye-field transcription factor *six3b* revealed substantial overlap, whereas rotation of the eGFP channel resulted in markedly reduced overlap. Together, these controls support the biological specificity of the observed overlap between *1cov*- and *3ov*-driven *eGFP* and endogenous *rax* transcripts during early eye development.

### Optic cup maturation

For additional evidence that these two enhancers drive expression strongly correlated with *rax*, enhancer-driven transgene expression and endogenous *rax* expression were assessed at 24 hpf. At this stage, endogenous *rax* transcripts were no longer detected within the optic cup, consistent with the previously described progressive loss of ocular *rax* expression as eye development proceeds. Similarly, *1cov*-driven expression was absent from the optic cup, although expression persisted within the hypothalamus, consistent with endogenous *rax* expression at this stage (Supplementary Fig. 2M-O). In contrast, for *3ov*-driven expression, persisted weakly within the optic cups, with the most prominent expression observed in the telencephalon, which is not shared with *rax* expression (Supplementary Fig. 2J-L).

Co-occurrence analysis using JaCop provides strong support for *1cov* and *3ov* being genuine *rax* enhancers. Both overlap substantially with endogenous *rax* transcripts during early ocular developmental stages. Additionally, *3ov*-driven expression changes in a manner similar to endogenous *rax* expression during transition from eye field specification to optic vesicle morphogenesis. For *1cov*, the expression dynamics are more strongly conserved with *rax,* as e*GFP* transcripts appear undetectable at 24 hpf, similar to *rax*.

### Association of 1cov-driven expression with *rax* transcripts at 72 hpf

At 72 hpf, 1cov drove strong expression within the hypothalamus, while retinal expression appeared localized to the INL (Fig. 3A). To assess the cell type specificity of *1cov* activity within the retina, cryosections were stained with anti-GFP antibodies. Labeled cells exhibited a characteristic radial morphology, with elongated processes spanning the entire thickness of the retina and cell bodies located within the INL, consistent with Müller glia cells (Fig. 3B)^49,50^.

At transcript level, co-occurrence analysis revealed limited overlap between 1cov-driven transcripts and endogenous *rax* transcripts, despite both signals being present within the INL (Fig. 3C-E). Endogenous *rax* transcripts were concentrated toward the outer edge of the INL and overlapped with 19% of 1cov driven transcripts (tM1 = 0.19 ± 0.05) (Fig. 3C). In contrast, the 1cov transcript signal was associated with cell bodies located within the INL and extended toward both the outer nuclear layer and ganglion cell layer, co-occurring with 29 % of *rax* transcripts (tM2 = 0.29 ± 0.09) (Fig. 3D). Thus, while both expression domains occupied the same retinal layer, their cellular or subcellular distributions appeared distinct. Despite these differences, the association between *rax* and eGFP was stronger than that observed in the negative control generated by rotating the eGFP channel 90° clockwise (tM1 = 0.18 ± 0.09; tM2 = 0.15 ± 0.13). *Rax* and *six3b* expression patterns diverge within the premature retina, although overlap is still shared among some retinal cell types. Consistent with this observation, thresholded Manders values demonstrated weaker shared expression between these two eye field transcription factors at this stage (Supplemental Fig. 3A-C). Notably, *eGFP* expression exhibited greater overlap with *rax* than *six3b.* Additionally, expression of the eye-specific gene *cplx4a* was examined because it is proximal to *rax*. *Cplx4a* is found within retinal ribbon synapses associated with photoreceptor and bipolar neurons in zebrafish^51^; however, its spatial expression pattern was not consistent with the distribution observed for the 1cov enhancer (Supplementary Fig. 3G-J).

In contrast to the retina, transcript co-occurrence analysis within the hypothalamus demonstrated substantial overlap between *rax*-positive and *eGFP*-positive pixels (Fig. 3F-H). A large proportion of *rax*-positive pixels overlapped with enhancer-driven expression (tM1 = 0.74 ± 0.03), whereas the proportion of eGFP-positive pixels overlapping with *rax* expression was comparatively lower (tM2 = 0.31 ± 0.05), reflecting the broader expression domain driven by the 1cov enhancer in this tissue (Fig. 3H). By comparison, the negative control, eGFP channel rotated 90° clockwise, revealed weaker co-occurrence signals (tM1 = 0.16 ± 0.10; tM2 = 0.12 ± 0.04). In the hypothalamus, thresholded Manders analysis revealed stronger overlap between *six3b* and *rax* than between *rax* and eGFP, although a larger proportion of *rax* transcripts overlap with eGFP transcripts (Fig. 3H; Supplemental Fig. 3D-F).

Together, these findings demonstrate that 1cov remains active at 72 hpf and continues to drive expression within domains associated with endogenous *rax* expression. While retinal transcript overlap is limited despite shared localization within the INL, hypothalamic expression exhibits substantial co-occurrence with endogenous *rax*. Furthermore, the elongated radial morphology and localization of cell bodies to the INL of 1cov-positive retinal cells are consistent with the morphology of Müller glia cells.

### Loss of 1cov enhancer destabilizes the *rax* GRN, resulting in ocular anomalies

To assess the developmental consequences of independently deleting each enhancer,*1cov* and *3ov*, from the genome, CRISPR-Cas9-mediated genome editing was employed. Guide RNAs were designed to target sequences flanking each enhancer, resulting in complete excision of the regulatory region (Fig. 4A; Supplementary Fig. 4A) Although no overt developmental abnormalities were observed in single enhancer deletion mutants, two standard eye measurements, axial eye and coronal diameter, normalized to larval standard length, were measured at protruding mouth stage (72 hpf) to identify more subtle effects on ocular development.

**Figure 4.**
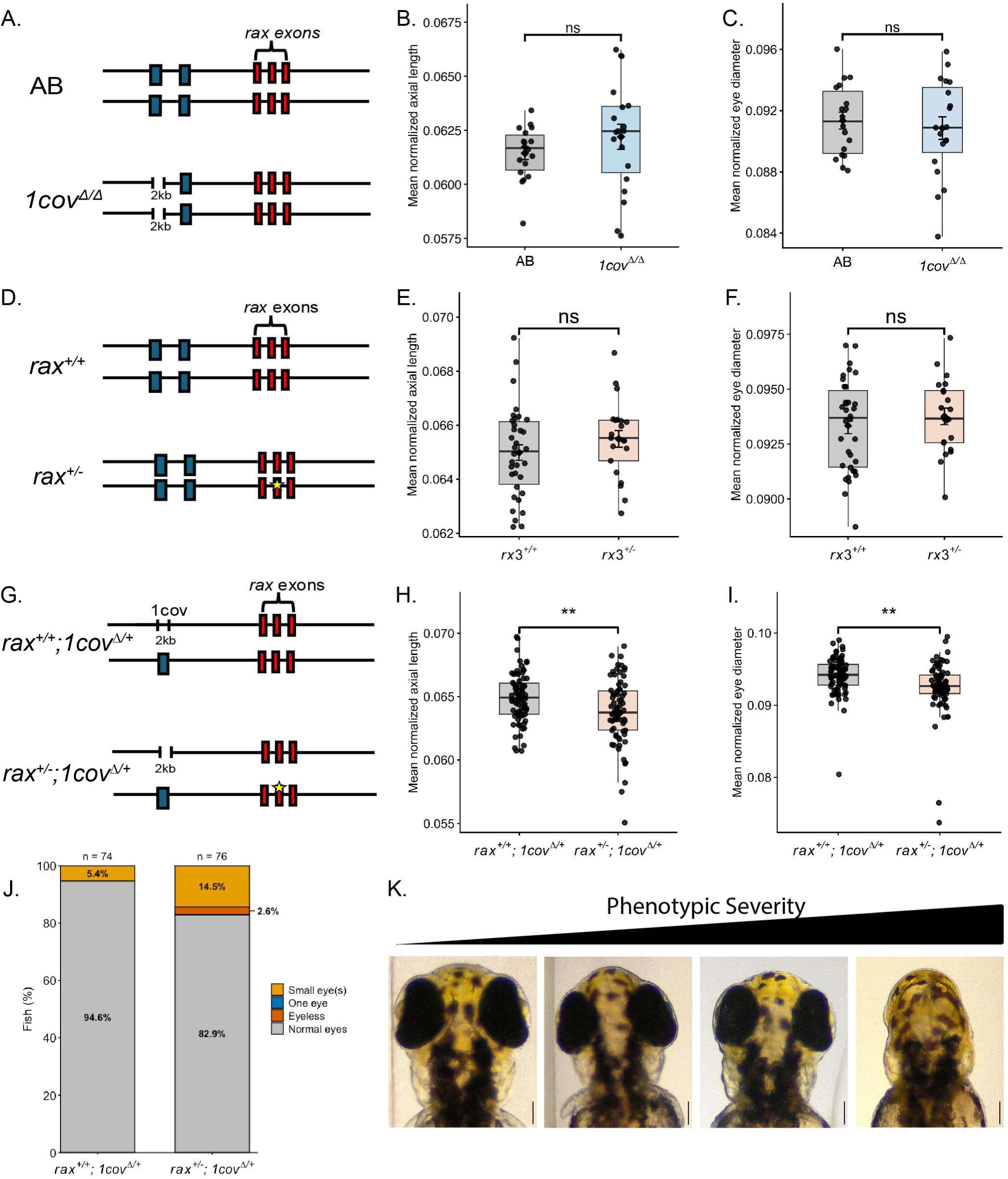
Loss of 1cov destabilizes *rax* regulation, leading to increased variance in eye size. (A) Schematic depicting the two genotypes compared in (B–C): AB wild-type and *1cov^Δ/Δ^*. (B) Comparison of normalized mean axial eye length between AB wild-type (*n* = 18) and *1cov^Δ/Δ^* (*n* = 19) larvae. Welch’s *t*-test, *p* = 0.25; F-test for equality of variances, *p* = 0.005. (C) Comparison of normalized mean coronal eye diameter between AB wild-type (*n* = 18) and *1cov^Δ/Δ^* (*n* = 19) larvae. Welch’s *t*-test, *p* = 0.60; F-test for equality of variances, *p* = 0.21. (D) Schematic depicting the two genotypes compared in (E–F): *rax*^+/+^ and *rax*^+/-^. The yellow star indicates a 12-bp insertion within *rax* exon 2. (E) Comparison of normalized mean axial eye length between *rax*^+/+^ (*n* = 34) and *rax*^+/-^ (*n* = 21) larvae. Welch’s *t*-test, *p* = 0.24; F-test for equality of variances, *p* = 0.45. (F) Comparison of normalized mean coronal eye diameter between *rax*^+/+^ (*n* = 34) and *rax*^+/-^ (*n* = 21) larvae. Welch’s *t*-test, *p* = 0.41; F-test for equality of variances, *p* = 0.35. (G) Schematic depicting the two genotypes compared in (H–J): *rax*^+/−^; *1cov^Δ/+^* and *rax*^+/+^; *1cov^Δ/+^* The yellow star indicates the 12-bp insertion within *rax* exon 2. (H) Comparison of normalized mean axial eye length between *rax*^+/-^; *1cov^Δ/+^* (*n* = 74) and *rax*^+/+^; *1cov^Δ/+^* (*n* = 74) larvae. Welch’s *t*-test, *p* = 0.005; F-test for equality of variances, *p* = 0.024. (I) Comparison of normalized mean coronal eye diameter between *rax*^+/-^; *1cov^Δ/+^* (*n* = 74) and *rax*^+/+^; *1cov^Δ/+^* (*n* = 74) larvae. Welch’s *t*-test, *p* = 0.002; F-test for equality of variances, *p* = 0.002. (J) Frequency of abnormal eye phenotypes in *rax*^+/-^; *1cov^Δ/+^* (*n* = 76) and *rax*^+/+^; *1cov^Δ/+^* (*n* = 74) larvae. Fisher’s exact test, *p* = 0.037. (K) Representative images depicting the range of severe ocular phenotypes observed among larvae from the cross shown in. Scale bars indicate 100µm. For (A–C), data were obtained from two independent clutches; for (D–F) and (G–I), data were obtained from three independent clutches. For quantitative eye-size measurements, only larvae with measurable bilateral eyes were included. Phenotypic frequency analyses included all larvae, including individuals with severe ocular phenotypes (anophtalmia) that could not be included in quantitative eye-size measurements.

Larvae homozygous for the *1cov* deletion (*1cov^Δ/Δ^*) exhibited significantly increased variance in normalized mean axial eye length compared with AB wild-type larvae (F-test, *p* = 0.005), although no significant differences in mean axial eye length or coronal diameter were observed between genotypes (Fig. 4B-C). When the two eyes were analyzed individually, the left eye exhibited significantly greater variance in axial eye length in *1cov^Δ/Δ^* larvae compared with AB wild-type larvae (F-test, *p* = 0.01), whereas the right eye did not, no other measurement were significant (Supplementary. Fig. 4C-D, F-G). In addition, no significant differences in axial eye length or coronal diameter were observed between AB wild-type and *3ov^Δ/Δ^* larvae (Supplementary Fig. 4C-G). Together, these observations suggest that loss of the *1cov* enhancer increases developmental variation in eye growth despite the absence of a consistent reduction in mean eye size.

To further examine the phenotypic variability observed following loss of *1cov*, we generated a sensitized genetic background by crossing fish carrying the homozygous *1cov* deletion with a fish heterozygous for a 12-bp loss-of-function allele within the DNA-binding domain of *rax*. Homozygosity for this 12-bp insertion allele results in a complete absence of eyes, and the location of the allele is indicated by a yellow star in the schematic (Fig. 4D; Supplementary Fig. 4H-I). This approach was designed to test whether reducing *rax* dosage while simultaneously removing regulatory input from *1cov* would produce a sensitized state in which the remaining functional copy of *rax* lacks regulatory input from this enhancer. To first determine whether heterozygous loss of *rax* function alone altered eye size, *rax*^+/-^ fish were crossed with AB wild-type fish, and axial eye length and coronal diameter were measured in their offspring. No significant differences were observed between *rax*^+/+^ and *rax*^+/-^ larvae (Fig. 4E-F; Supplementary Fig. 4J-M), consistent with previous observations in *rax* heterozygous loss-of-function fish in *Astyanax mexicanus*^45^.

Following validation of this experimental design, *1cov^Δ/Δ^* fish were crossed with *rax*^+/-^ fish, and normalized axial eye length and coronal diameter were measured at the protruding-mouth stage in sensitized double-heterozygous larvae and compared with control siblings carrying two wild-type copies of *rax* and a heterozygous deletion of the *1cov* enhancer (Fig. 4G). Sensitized *rax*^+/-^; *1cov^Δ/+^* larvae exhibited significantly reduced mean normalized axial eye length compared with *rax*^+/+^; *1cov^Δ/+^* siblings (Welch’s *t*-test, *p* = 0.005) and significantly increased variance (F-test, *p* = 0.024) (Fig. 4H). Similarly, sensitized larvae exhibited significantly reduced mean normalized coronal diameter (Welch’s *t*-test, *p* = 0.002) and significantly increased variance (F-test, *p* = 0.0002) (Fig. 4I). As a conservative measure, anophthalmic larvae, which only occurred in the sensitized *rax^+/-^; 1cov^Δ/+^* genotype, were excluded from these measurements. When individual eyes were analyzed separately, significant differences were observed for all comparisons except variance in the axial eye length of the left eye (Supplementary Fig. 4N-Q).

Affected larvae were identified by first calculating the mean of the left and right eye measurements for each individual *rax***^+/+^;** *1cov^Δ/+^* control larva. These individual mean eye measurements were then used to calculate the group mean and standard deviation for each ocular measurement. A threshold was established at two standard deviations below the group mean of rax^+/+^ sibling controls for both normalized axial eye length and coronal diameter, and any eye measurement falling below either threshold was classified as microphthalmic^52,53^. Larvae with one or both eyes below this threshold were considered affected, with each larva counted as a single affected individual. Within the *rax*^+/-^; *1cov^Δ/+^* genotype, thirteen larvae exhibited microphthalmic phenotypes and two exhibited bilateral anophthalmia. In comparison, four *rax*^+/+^; *1cov^Δ/+^* larvae exhibited microphthalmic phenotypes (Fig. 4J). Thus, the sensitized genotype was enriched for larvae with small eyes and included two larvae with the severe ocular phenotype of bilateral anophthalmia. The association between the sensitized genotype and abnormal ocular phenotypes was significant (Fisher’s exact test, *p* = 0.027). Overall, larvae from the sensitized background exhibited a range of phenotypic severity, from normal eyes to anophthalmia, although most affected individuals exhibited relatively mild phenotypes (Fig. 4K).

Despite the limited penetrance of overt ocular phenotypes, larvae in the sensitized population had smaller eyes overall and greater variation in eye size, with the range and severity of phenotypic outcomes exceeding those observed in the homozygous *1cov* deletion background alone. These findings support the notion that loss of *1cov* destabilizes the *rax* GRN, increasing developmental variability and likely sensitizing ocular development to additional genetic perturbations.

### Progressive regulatory perturbation increases the severity and range of ocular phenotypes

Given the role of *1cov* in supporting *rax* regulation during eye development, we next used CRISPR/Cas9 to target the *3ov* enhancer for deletion in the sensitized genetic background. We hypothesized that, if *3ov* functions as a *rax* enhancer, its deletion would further disrupt *rax* regulation, resulting in increased penetrance and severity of ocular phenotypes in sensitized *rax*^+/-^; *1cov^Δ/+^* larvae (Fig. 5G). We also performed the reciprocal experiment, in which fish harboring a homozygous deletion of the *3ov* enhancer were crossed with *rax*^+/-^ fish, and the resulting embryos were injected with guides targeting the *1cov* enhancer for deletion (Fig. 5D). Thus, the two experiments examined secondary enhancer loss in both directions, with *3ov* targeted in the *1cov* deletion background and *1cov* targeted in the *3ov* deletion background. Importantly, unlike the homozygous *1cov* deletion background, fish homozygous for the *3ov* deletion did not display overt differences in eye size when crossed with *rax*^+/-^ fish. As a negative control, we also crossed *1cov* homozygous deletion fish with *rax*^+/-^ fish and injected the resulting embryos with a guide targeting eGFP (Fig. 5A)^54^. This control was used to determine whether the injection procedure itself contributed to the ocular phenotypes observed following CRISPR/Cas9-mediated targeting of the secondary enhancers.

**Figure 5.**
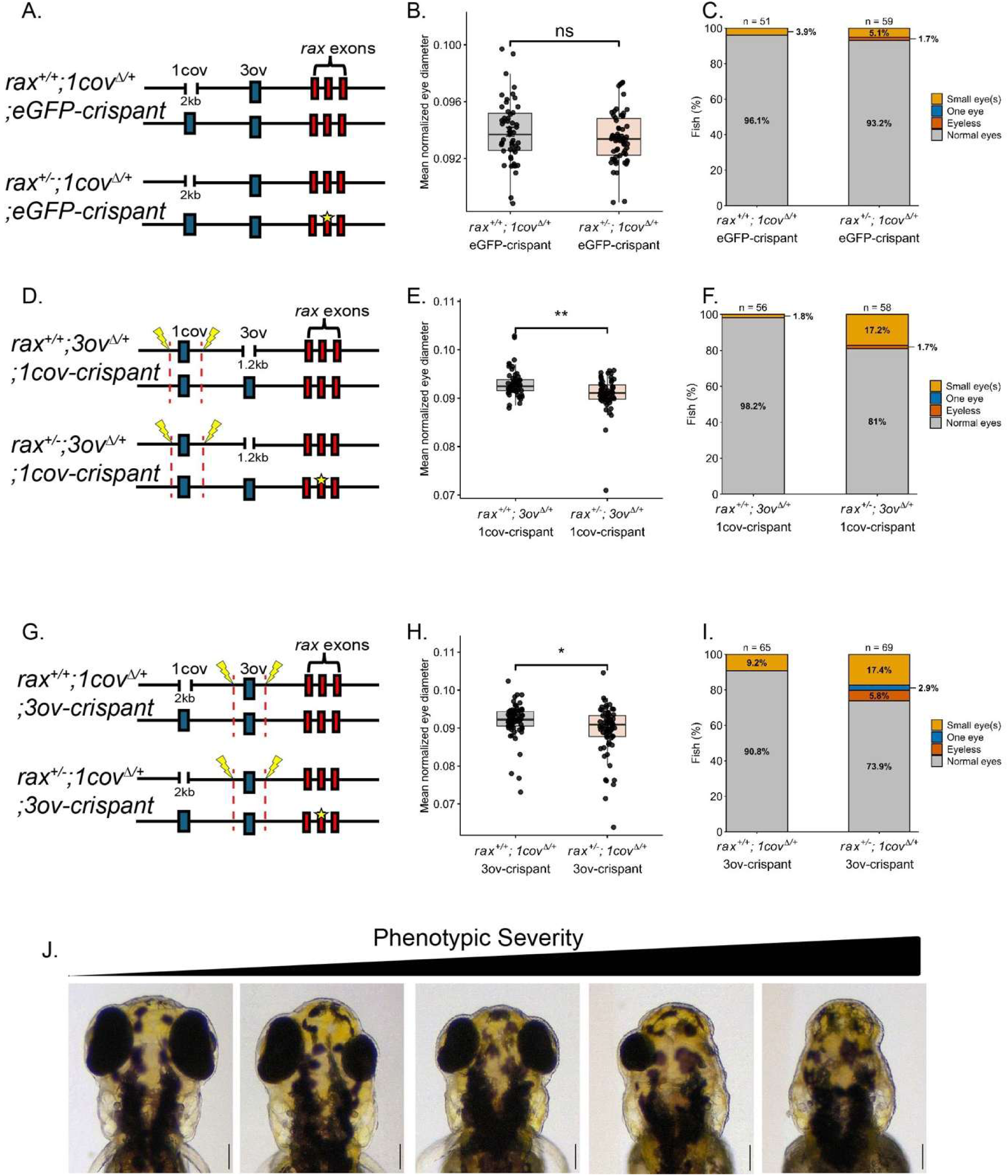
Genomic excision of secondary *rax* enhancers increases phenotypic severity. (A) Schematic depicting the two genotypes resulting from a cross between *1cov^Δ/Δ^* and *rax*^+/-^ fish. Embryos were injected with guides targeting eGFP as a control for injection-associated phenotypes. (B) Boxplots depicting mean normalized coronal eye diameter of larvae from the two genotypes shown in (A): *rax*^+/+^; *1cov^Δ/+^*; eGFP-crispant (*n* = 51) and *rax*^+/-^; *1cov^Δ/+^*^;^ eGFP-crispant (*n* = 59). Welch’s *t*-test, *p* = 0.27. (C) Percentage of larvae from the cross shown in (A) exhibiting normal eyes, small eye(s), one eye, or no eyes: *rax*^+/+^; *1cov^Δ/+^;* eGFP-crispant (*n* = 51) and *rax*^+/-^; *1cov^Δ/+;^* eGFP-crispant (*n* = 60). Fisher’s exact test, *p* = 0.684. (D) Schematic depicting the two genotypes resulting from a cross between *3ov^Δ/Δ^* and *rax*^+/-^ fish. Embryos were injected with guides targeting the 1cov enhancer for genomic excision. (E) Boxplots depicting mean normalized coronal eye diameter of larvae from the two genotypes shown in (D): *rax^+/+^; 3ov^Δ/+^* ;1cov-crispant (*n* = 56) and *rax^+/-^; 3ov^Δ/+^*; 1cov-crispant *(n* = 57). Welch’s *t*-test, *p* = 0.001.(F) Percentage of larvae from the cross shown in (D) exhibiting normal eyes, small eye(s), one eye, or no eyes: *rax^+/+^; 3ov^Δ/+^*; 1cov-crispant *(n* = 56) and *rax^+/-^; 3ov^Δ/+^*; 1cov-crispant (*n* = 58). Fisher’s exact test, *p* = 0.004. (G) Schematic depicting the two genotypes resulting from a cross between 1cov^Δ/Δ^ and *rax*^+/-^ fish. Embryos were injected with guides targeting the *3ov* enhancer for genomic excision. (H) Boxplots depicting mean normalized coronal eye diameter of larvae from the two genotypes shown in (G): *rax*^+/+^; *1cov^Δ/+;^* 3ov-crispant (*n* = 65) and *rax*^+/-^; *1cov^Δ/+^*^;^ 3ov-crispant (*n* = 69). Welch’s *t*-test, *p* = 0.015. (I) Percentage of larvae from the cross shown in (G) exhibiting normal eyes, small eye(s), one eye, or no eyes: *rax*^+/+^; *1cov^Δ/+^*; 3ov-crispant (*n* = 65) and *rax*^+/-^; *1cov^Δ/+^*^;^ 3ov-crispant (*n* = 73). Fisher’s exact test, *p* = 0.013. (J) Representative images depicting the range of eye phenotype severity observed in larvae from the cross shown in. Scale bars indicate 100µm. For all experiments, the reported *n* values represent the total number of larvae pooled across three independent clutches. Quantitative eye-size analyses included larvae with at least one measurable eye, whereas phenotypic frequency analyses included all larvae, including individuals with bilateral anophthalmia that could not be included in quantitative eye-size measurements.

In these experiments, normalized coronal diameter more clearly captured the phenotypic variability observed between genotypes and was therefore used as the primary quantitative measure in Fig. 5. Some larvae displayed normalized axial eye lengths within the range observed for normal eyes while exhibiting substantially reduced coronal diameters characteristic of severe microphthalmia. Axial eye length measurements are provided in Supplementary Fig. 5.

In the eGFP-injected negative control, the differences between genotypes observed in the previous cross (Fig. 4H-J) were not detected. No significant difference in mean normalized coronal diameter was observed between *rax*^+/-^; *1cov^Δ/+^* and *rax*^+/+^; *1cov^Δ/+^* larvae (Welch’s *t*-test, *p* = 0.27), and no significant difference in variance was detected (F-test, *p* = 0.15) (Fig. 5B). Individual coronal diameter measurements of the left and right eyes were consistent with the mean eye measurements (Supplementary Fig. 5A-B). Although the difference in mean normalized axial eye length approached statistical significance, only the left-eye axial length differed significantly between genotypes, whereas the right-eye axial length did not (Supplementary Fig. 5C-E). Furthermore, the sensitized genotype was not enriched for ocular phenotypes in the eGFP-injected control (Fig. 5C; Fisher’s exact test, *p* =0.68).

To test whether loss of *1cov* could produce notable ocular phenotypes in the *3ov* deletion background, *3ov^Δ/Δ^* fish were crossed with *rax*^+/-^ fish, producing *rax^+/+^; 3ov^Δ/+^* and sensitized *rax*^+/-^; *3ov^Δ/+^* larvae. The resulting embryos were injected with guides targeting the *1cov* enhancer for deletion (Fig. 5D). Excluding anophthalmic larvae, normalized coronal diameter was compared between the two genotypes. Sensitized *rax*^+/-^; *3ov^Δ/+^* larvae exhibited a significantly reduced mean normalized coronal diameter compared with *rax*^+/+^; *3ov^Δ/+^* larvae (Welch’s *t*-test, *p* = 0.001), although the difference in variance did not reach statistical significance (F-test, *p* = 0.058) (Fig. 5E). In this experiment, affected larvae where highly enriched in sensitized background, with 18.9% of *rax*^+/-^; *3ov^Δ/+^* larvae affected compared with 1.8% of *rax*^+/+^; *3ov^Δ/+^* larvae (Fig. 5F; Fisher’s exact test, p= 0.004). Individual analysis of the left and right eyes similarly revealed significant differences in normalized coronal diameter between genotypes, with significantly greater variance observed for the left eye but not the right eye (Supplementary Fig. 5F-G). In contrast, normalized axial eye length did not differ significantly between the two genotypes (Supplementary Fig. 5H-J).

In a similar cross using the genotypes depicted in Fig. 4G, 1cov^Δ/Δ^ fish were crossed with *rax*^+/-^ fish, producing *rax*^+/+^; *1cov^Δ/+^* and sensitized *rax*^+/-^; *1cov^Δ/+^* larvae. The resulting embryos were injected with guides targeting the *3ov* enhancer for deletion (Fig. 5G). Excluding anophthalmic larvae, normalized coronal diameter measurements revealed a significant difference in mean eye size between genotypes (Welch’s *t*-test, *p* = 0.015), with significantly greater variance in the sensitized genotype (F-test, *p* = 0.004) (Fig. 5H). Significant differences in normalized coronal diameter were also observed when the left and right eyes were analyzed individually, along with significantly higher variance (Supplementary Fig. 5K-L). In contrast, normalized axial eye length did not differ significantly between genotypes, although a significant difference in variance was observed for the right eye (Supplementary Fig. 5M-O). The sensitized *rax*^+/-^; *1cov^Δ/+^* genotype exhibited affected ocular phenotypes in 26.1% of larvae, compared with 9.2% of *rax*^+/+^; *1cov^Δ/+^* larvae (Fig. 5I). Fisher’s exact test demonstrated a significant association between the sensitized genotype and affected ocular phenotypes (*p* = 0.013). The ocular phenotypes observed in this cross encompassed a broad range, from normal eyes to complete bilateral anophthalmia, with greater phenotypic diversity than observed in the other crosses (Fig. 5J).

Together, although the phenotypic differences observed in non-injected larvae were not detected in the eGFP-injected control, targeted deletion of either secondary enhancer consistently resulted in ocular phenotypes that were enriched in the sensitized background. Phenotypic variability and penetrance were greatest when the *1cov* enhancer was already absent from the functional *rax* allele, and the *3ov* enhancer was subsequently targeted for deletion. Ocular phenotypes were also observed in the reciprocal experiment, in which the *3ov* enhancer was deleted and *1cov* was subsequently targeted in the *rax* sensitized background. Collectively, these results demonstrate that perturbation of *1cov* and *3ov* increases the range and penetrance of ocular phenotypes in a *rax* sensitized background.

## Discussion

### Characterization of the *rax* enhancer landscape

Together, these findings identify five previously uncharacterized cis-regulatory elements associated with *rax* and demonstrate that each drives expression within embryonic ocular tissues where *rax* is normally expressed. The shared activity of multiple enhancers within *rax*-associated domains suggests that *rax* regulation is governed by a broader cis-regulatory landscape than previously appreciated. Detailed characterization of *1cov* and *3ov* revealed substantial overlap between enhancer-driven reporter activity and endogenous *rax* transcripts, supporting their role as biologically relevant components of the *rax* regulatory network. Importantly, both enhancers largely recapitulated endogenous *rax* expression domains, with differences arising primarily from additional reporter activity in adjacent tissues rather than a failure to reproduce known *rax* expression territories. Such discrepancies are common in transgenic enhancer assays and likely reflect the removal of enhancers from their endogenous regulatory context, where interactions with additional activating and repressive cis-regulatory inputs, and endogenous promoter contribute to refining spatial and temporal expression patterns ^55–57^.

The shared retinal and hypothalamic expression observed by some reporter constructs further suggests that *rax* expression is not partitioned among discrete enhancers controlling independent developmental domains. Rather, multiple enhancers drive overlapping expression within shared ocular territories, consistent with a distributed regulatory architecture^36,58^. Such architecture could provide robustness to this developmental expression program. While embryonic ocular expression was observed across all five enhancers, later hypothalamic activity appeared unique to *1cov*, suggesting that individual enhancers may also contribute specialized regulatory inputs at specific developmental stages or within particular cell populations^31^.

### Spatial and temporal complexity of *rax* enhancer activity

At later developmental stages, colocalization between *1cov* activity and endogenous *rax* expression within the retina was reduced relative to earlier stages. However, this decrease likely reflects the highly restricted localization of endogenous *rax* transcripts to the outer edge of the INL rather than a complete divergence of enhancer activity from endogenous gene expression. If *rax* were expressed uniformly throughout bipolar cells, transcript signal would be expected to span a greater proportion of the INL. Instead, the preferential localization of *rax* transcripts to the outer region of the INL suggests that subcellular or cell-type-specific transcript localization contributes to the weak co-occurrence values observed at protruding mouth stage despite both expression domains being present within the same retinal layer. Spatially restricted transcript localization is well established in neural and glial cells, where specific mRNAs can be transported to and enriched within distinct subcellular regions^59,60^.

Notably, *1cov* activity was observed within Müller glia, a finding that is biologically plausible given reports of *Rax* activity in analogous retinal cell populations in mammals^9,22^. Conversely, if *rax* expression within bipolar cells is regulated by distinct cis-regulatory elements, such enhancers may not have been recovered by the workflow used in this study and would require more informative datasets that align with the onset of bipolar cell differentiation. *Rax* is expressed within the hypothalamus at this stage, which was recapitulated by *1cov* transgene expression, albeit with a somewhat broader expression domain. The broader hypothalamic expression of *1cov* is consistent with the established role of *rax* in the developing zebrafish hypothalamus, where *rax* marks and regulates tuberal/anterior hypothalamic progenitors^10^. However, the broader distribution of *1cov* activity relative to endogenous *rax* expression may reflect the removal of this enhancer from its native cis-regulatory context.

The overlapping activities observed across multiple enhancers indicate that *rax* expression is robustly regulated during eye field specification and optic vesicle evagination, a developmental period in which *rax* function is critical for normal optic vesicle morphogenesis^3^. Importantly, this apparent reduction in enhancer activity at later stages may reflect a bias in candidate prioritization, as optimal chromatin-based datasets were not available for later timepoints. These observations suggest that *rax* expression is controlled by a dynamic enhancer landscape in which distinct regulatory elements contribute during different developmental windows, during early ocular development. However, at later timepoints, when *rax* plays a more nuanced role in the developing eye, *rax* is likely regulated by distinct enhancers directing expression within the retinal subtypes. Such an organization is consistent with enhancer redundancy and likely contributes to the robust maintenance of *rax*-dependent developmental programs^37,58^.

### Role of *1cov* in developmental robustness

Deep functional analysis of two novel ocular enhancers, *1cov* and *3ov*, supports the existence of a robust GRN underlying *rax* expression. Although this regulatory network is resilient to perturbation, it nevertheless appears to require input from specific enhancers during key developmental windows, particularly during optic vesicle evagination and early proliferative stages.

We propose that the deeply conserved *1cov* enhancer functions as a buffering element that contributes to the stability of *rax* dependent ocular development during these periods. While genetic ablation of *1cov* alone is largely tolerated under standard lab conditions, similar tolerance of individual enhancer loss has been observed in other developmental contexts, with enhancer requirements becoming more apparent when target gene dosage is reduced and with individual enhancer loss being associated with increased phenotypic variability^37,61,62^. Consistent with this model, the consequences of *1cov* loss became more apparent when *rax* dosage was reduced, with sensitized individuals exhibiting microphthalmic and, less frequently, anophthalmic phenotypes. These findings suggest that the contribution of *1cov* to ocular development is dependent on the broader regulatory architecture acting at the *rax* locus.

### Progressive regulatory perturbation increases frequency and range of ocular phenotypes

Although the reduction in eye size and increased phenotypic variability observed in the sensitized *1cov* background were not maintained following control injection with eGFP guides, targeted deletion of a second enhancer produced more consistent phenotypic differences. Targeting *3ov* in the stable *1cov* deletion background resulted in reduced coronal eye diameter, increased penetrance, and phenotypic variability in sensitized larvae. Similarly, targeting *1cov* in the stable *3ov* deletion background recapitulated ocular abnormalities in the sensitized genotype. Together, these reciprocal perturbations provide genetic evidence that both *1cov* and *3ov* contribute to *rax* regulation and ocular development. Combinatorial dependencies among cis-regulatory elements have also been observed in the regulation of other genes, where combined perturbation can reveal regulatory requirements that are less apparent following disruption of a single enhancer^36,37^.

Despite progressive disruption of the *rax* regulatory landscape, the majority of sensitized larvae developed normal eyes, suggesting that substantial regulatory buffering persists. This incomplete penetrance may reflect regulatory activity from the *rax* promoter and additional cis-regulatory elements. Nevertheless, the increased variability observed following loss of *1cov*, together with the broader range and increased penetrance of phenotypes following perturbation of an additional enhancer, supports a role for *1cov* in maintaining the robustness of the *rax* regulatory network.

The range of phenotypes observed following progressive regulatory perturbation may also reflect disruption of *rax* regulation at different stages of eye development. Severe phenotypes, including anophthalmia and pronounced microphthalmia, may result from disruption of *rax* regulation during early optic vesicle evagination, whereas milder microphthalmic phenotypes may reflect perturbation of later processes contributing to retinal progenitor proliferation and ocular growth^3,63,64^. Collectively, these findings support a model in which ocular defects emerge through the progressive erosion of regulatory robustness across the *rax* locus rather than through the loss of any single enhancer.

## Methods

### Zebrafish Husbandry

Zebrafish embryos were collected and stored at 28.5 °C in either E3 embryo media or E3 containing 0.003% 1-phenyl-2-thiourea when pigmentation inhibition was necessary to maintain transparency at later developmental stages. Embryos were staged according to morphological criteria and collected at the appropriate developmental stage for each experiment^48^. Prior to experimental analysis, larvae were euthanized by immersion in 0.3g/L MS-222 (Syncaine) 30 minutes.

All zebrafish experiments were approved by the University of Pittsburgh Institutional Animal Care and Use Committee (IACUC) and performed in accordance with approved protocols.

### Putative Enhancer selection

Candidate regions were selected based on one or more of the following criteria: proximity to *rax* within a two-megabase window, evidence of H3K27ac enrichment, evolutionary conservation, and detectable chromatin accessibility in the optic vesicles. Publicly available datasets were used for enhancer selection. Chromatin accessibility (ATAC-seq) datasets generated in Prim-5 stage zebrafish were used to prioritize candidate regions with open chromatin in the optic vesicles^65^. ChIP-seq datasets generated from bud, 5-9 somite, and Prim-5 stage zebrafish were used to identify regions enriched for enhancer-associated histone marks H3K27ac and H3K4me1^66,67^. Lastly, 4-way (zebrafish, grass carp, common carp, and goldfish) PhastCons scores were used to evaluate conservation at these candidate regions, accessed through DANIO-CODE^68,69^.

### Generation of Expression Vector constructs

Candidate regions were amplified from AB-strain genomic DNA using Platinum SuperFi II DNA polymerase (Thermo Fisher, cat # 12361010). Standard thermocycling conditions were used, with annealing temperature and elongation time optimized for each region.

The five candidate regions, designated *1cov*, *2ov*, *3ov*, *4ov*, and *5ov* were cloned into enhancer-reporter constructs using either pEXPR:SA-E1B-egfp-pA-CR3(*1cov* and *3ov*) or pEXPR-FAbas-egfp-pA-CG2(*2ov*, *4ov*, and *5ov*). Cloning was performed using NEBuilder HiFi DNA Assembly master mix (NEB cat#E2621L).

The pEXPR-FAbas-egfp-pA-CG2 was digested with AscI (NEB, cat # R00558S) and FseI(NEB, cat # R0588S) using rcutsmart buffer and incubated at 37 °C for overnight. Inserts amplified with complementary overhangs matching the plasmid insertion sites were incorporated into the digested vector.

The pEXPR:SA-E1B-egfp-pA-CR3 vector was digested with ApaI (NEB, cat # R0114S) and SalI-HF (NEB, cat# R3138S) using rcutsmart buffer and incubated overnight at 37 °C. Inserts amplified using primers containing complementary overhangs were then assembled into the plasmid. Both expression vectors contain Tol2 recombination sites, enabling random genomic integration.

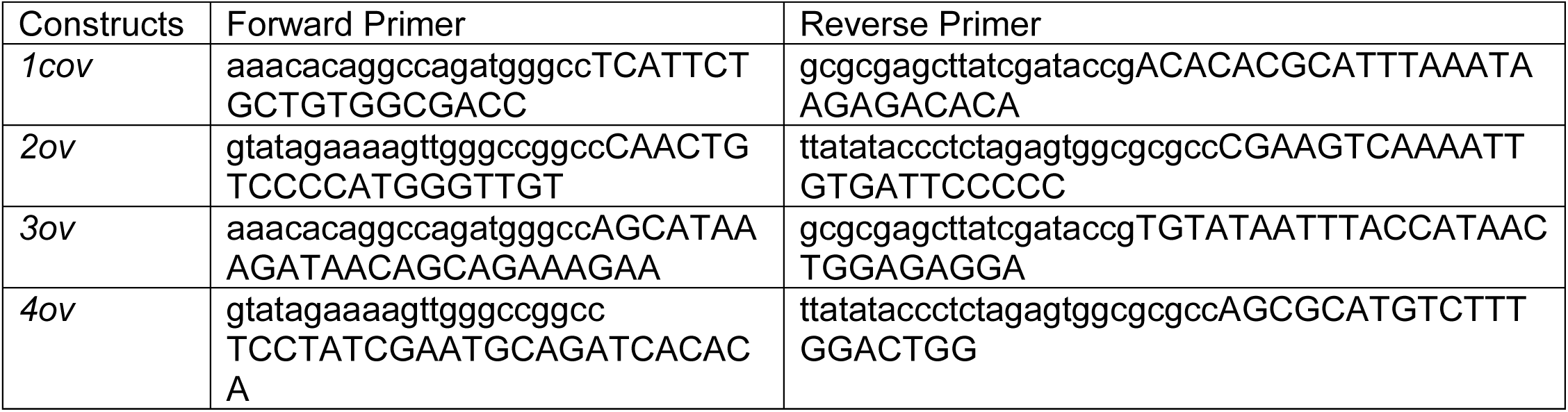
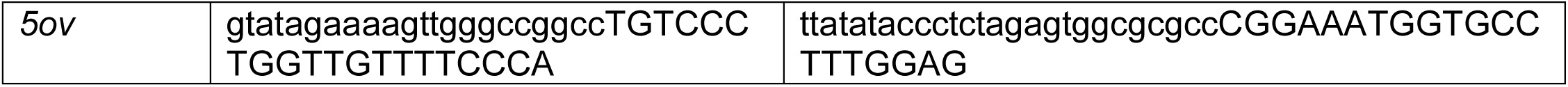

Assembled plasmids were then transformed into DH5-alpha chemically competent E. coli (Thermo Fisher cat# EC0112) by heat shock for 30 seconds at 42°C, and ampicillin-resistant colonies were selected. Plasmids were isolated by miniprep (Qiagen, cat #27106) and sequenced using long-read sequencing (Plasmidsaurus) to confirm the presence of the target region.

### Generation of Transgenic Lines

One nanoliter of a 10-µL injection cocktail was microinjected into AB-strain zebrafish embryos at the one-cell stage. The injection mix contained phenol red (1 µL), plasmid construct (35 ng/µL), Tol2 transposase mRNA (30 ng/µL), 0.4 M KCl (5 µL), and nuclease-free water to a final volume of 10 µL^70^. To control for positional effects resulting from random Tol2-mediated integration, expression patterns were determined by screening at least three independent founders per construct. Only F1 progeny displaying reproducible spatial and temporal expression patterns, as visualized by the fluorescent reporter at 24 and 48 hpf, were selected for line establishment.

### Live Transgenic Embryo Imaging

H2A-mCherry mRNA was synthesized *in vitro* from the pME-H2A-mCherry plasmid using the mMESSAGE mMACHINE SP6 Transcription Kit (Thermo Fisher Scientific) according to the manufacturer’s instructions. A working injection solution was prepared by diluting H2A-mCherry mRNA to a final concentration of approximately 104 ng/µL in nuclease-free water containing phenol red. One to two nanoliters of the injection mixture were injected into one-cell-stage embryos to visualize nuclei during live imaging.

### Generation of Enhancer Deletion Lines and *rax* Mutant Line

Enhancer deletion lines were generated using CRISPR-Cas9-mediated genome editing as previously described.^71^ Guide crRNAs targeting sequences flanking each enhancer were designed using the IDT CRISPR design tool^72^. crRNAs were annealed to tracrRNA to generate guide RNA (gRNA) duplexes. Ribonucleoprotein (RNP) complexes were assembled by incubating gRNA duplexes with Cas9 protein (IDT, cat. #10007806) for 5 min at 37°C. The injection cocktail consisted of 1 µL of 25 µM guide RNA 1, 1 µL of 25 µM guide RNA 2, 1 µL of 25 µM Cas9 protein, 1 µL phenol red, and 1 µL nuclease-free water. One nanoliter of the injection mixture was injected into one-cell-stage embryos.

The *rax* mutant line was generated using CRISPR-Cas9 genome editing targeting the DNA-binding domain within exon 2 of *rax*. Anophthalmic founder fish were crossed to wild-type animals by in vitro fertilization (IVF) and resulting F1 progeny were raised to adulthood^73^. Heterozygous carriers were subsequently intercrossed, producing offspring that exhibited the characteristic *rax* anophthalmic phenotype, at the proper mendelian ratios (∼25%). Sequencing of affected individuals identified a 12-bp insertion within the DNA-binding domain of exon 2. The allele was maintained as a heterozygous stock and used throughout this study.

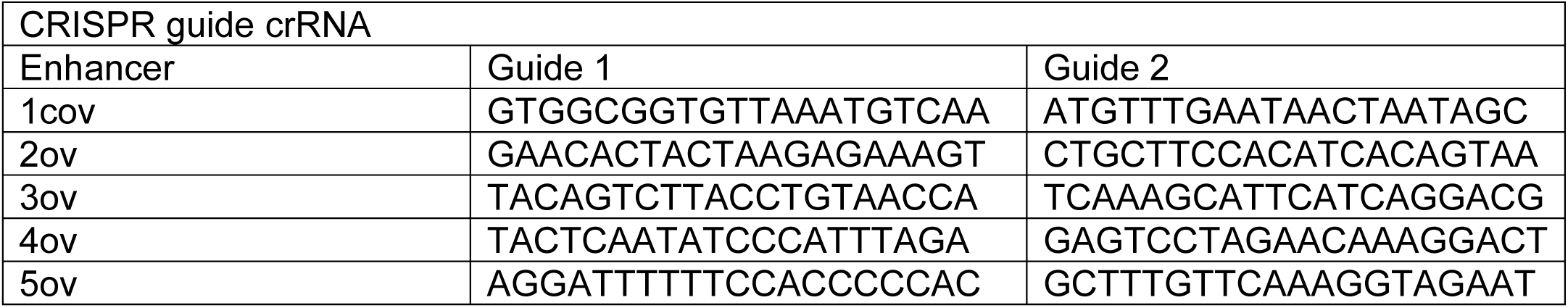
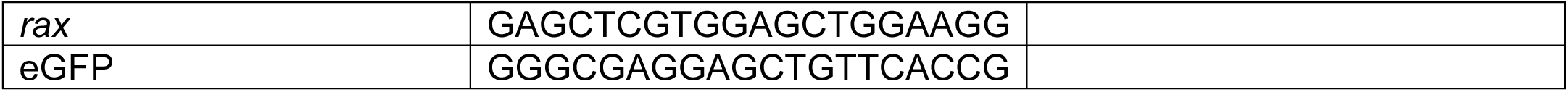

### Genotyping Embryos

Genomic DNA was extracted from embryos by incubating them in 50 µl of 50mM NaOH AT 95°C for 30 min, followed by the addition of 5 µl 1 M Tris, pH 8. Embryos were genotyped by PCR using primers flanking the gRNA target sites. Successful deletion events produced a substantially smaller amplicon relative to the wild-type allele. PCR products corresponding to deletion alleles were validated by Sanger sequencing. For rax^+/-^ larvae containing the loss of function allele was determined by PCR amplifying the region followed by restriction digestion with Fnu4HI for 1 hour at 37°C. The Fnu4HI restriction site was present in the mutant allele resulting in two fragments following digestion, visible by gel electrophoresis.

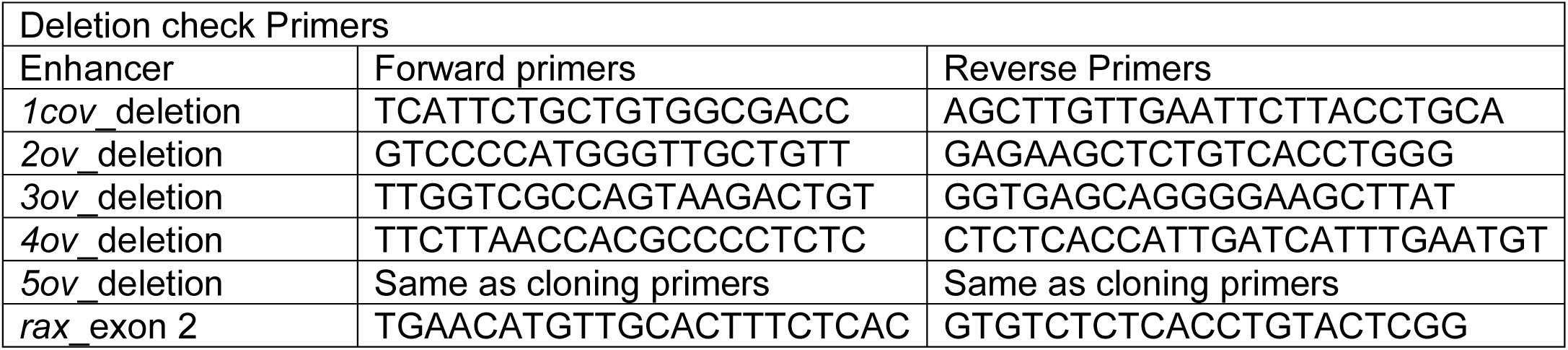

### Whole mount Hybridization chain Reaction (HCR)

HCR reagents, RNA-specific probes, and fluorophore-conjugated hairpins were acquired through Molecular instruments. Prior to HCR, embryos were fixed in 4% paraformaldehyde (PFA) for two hours at room temperature, washed in PBS and dehydrated in 100% methanol, and stored at -20 °C until use. HCR was performed according to manufacturer’s protocol provided by Molecular Instruments^74^.

| HCR probes used in this study |  |
|---|---|
| *rax*-x3 | Designed by Molecular Instruments |
| *eGFP*-x2 |  |
| *cplx4a*-x1 |  |
| *six3b*-x1 |  |

### Immunohistochemistry

Larvae used for cryosectioning were fixed overnight in 4% paraformaldehyde (PFA) at 4°C and washed three times in PBS. Samples were subsequently incubated overnight in 30% sucrose, embedded in OCT compound, and stored at −80°C until sectioning. Larvae were cryosectioned at 8 µm thickness.

Sections were permeabilized in 0.2% Triton X-100 for 10 min, blocked in 5% goat serum for 1 h, and incubated overnight at 4°C with anti-GFP antibody (Thermo Fisher, PA1-9533) at a dilution of 1:400 in blocking solution. Sections were then washed four times for 15 min in 0.2% PBST and incubated with goat anti-chicken Alexa Fluor 488 secondary antibody (Thermo Fisher, A11039) at a dilution of 1:1000 for 1 h. DAPI was included at a final dilution of 1:1000. Following secondary antibody incubation, sections were washed four times for 15 min in 0.1% PBST, followed by two 10 min washes in PBS, and mounted in glycerol for imaging.

### Image acquisition and Measurement

#### Confocal Imaging

Images were acquired using a Nikon A1R confocal microscope with 10x, 20x, and 40x objectives. Embryos were embedded in 1% low-melt agarose (Invitrogen) and imaged using 0.5-3 µm z-steps. Image analysis was performed using z-stack image series acquired through the specimen.

#### Stereo scope

Images were acquired at 4x magnification on Leica S9i. Tricaine euthanized larvae were place on laterally on a 6-lane injection mold to acquire images

#### Image Analysis

Colocalization analysis was performed in FIJI (ImageJ) using the JaCoP plugin on three independent images. Colocalization was quantified using thresholded Manders’ overlap coefficients (tM1 and tM2). Prior to analysis, a Gaussian blur (σ = 2) was applied to both channels to reduce background signal. Thresholds were manually set at the point where the primary observable fluorescence signal could be distinguished from background fluorescence before calculation of the colocalization coefficients. To provide biological context for the observed co-occurrence between enhancer-driven eGFP and endogenous *rax* transcripts, *six3b* was used as a positive control based on its overlapping expression with *rax* in anterior neural and ocular territories during early zebrafish development^75–77^, while the eGFP channel rotated 90° clockwise was used as a negative control.

#### Eye Measurement

Axial lengths were determined by measuring the distance from the tip of the cornea to the posterior edge of the retina using FIJI (ImageJ). Both eyes were measured in all embryos, and measurements from each eye were recorded independently and normalized to total body length^78^. Image acquisition, data collection, and data processing were performed prior to genotyping.

#### Scoring abnormal phenotypes

Homozygous mutant larvae (*1cov^Δ/Δ^* and *3ov^Δ/Δ^*) were embedded in 1% low-melt agarose and imaged using a confocal microscope. Following image acquisition, eye axial lengths were measured in FIJI and normalized to total larval length. Measurements were then compared to those obtained from wild-type (AB) siblings.

To evaluate the effects of a sensitized genetic background, rax^+/-^ zebrafish were crossed with homozygous enhancer deletion (*1cov^Δ/Δ^* and *3ov^Δ/Δ^*) zebrafish. Embryos were imaged at 72 hpf at 4× magnification using a Leica dissecting microscope. Axial length and coronal diameter (nasal to temporal length of eye) were measured prior to genotyping of the inherited *rax* allele, and normalized to the standard length of the larva ^78,79^. Embryos with one or both eyes exhibiting an axial length or coronal diameter two standard deviations below the population mean of both eyes from rax^+/+^ siblings were classified as microphthalmic. Embryos lacking one or both eyes were classified as anophthalmic and automatically classified as affected.

## Data Availability

All data generated or analyzed during this study are included in the article and its Supplementary Data files. Raw data for JaCoP co-occurrence analyses are provided in Supplementary Data Table 1. Raw axial eye length and coronal diameter measurements from 3 dpf zebrafish are provided in Supplementary Data Table 2. Representative microscopy images and additional source data supporting the findings of this study are available from the corresponding author upon reasonable request.

## Acknowledgements

We thank Dr. Kristen Kwan (University of Utah) for providing the or pCS2FA-H2A.F/Z-mCherry plasmid and advice provided for this project.

## Use of Generative AI tools

Claude (Sonnet 5, Anthropic) and Microsoft Copilot were used solely to generate scripts for data analysis and visualization in R Studio, and to provide editorial assistance for improving readability. All scientific content was written reviewed and verified by the authors.

